# Similarity bias from consensus perturbational signatures from the L1000 Connectivity Map

**DOI:** 10.1101/2022.01.24.477615

**Authors:** Ian Smith, Katy Scott, Benjamin Haibe-Kains

## Abstract

In recent years, high-throughput perturbational datasets have become an important tool for rapidly characterizing the function of large collections of chemical compounds. To overcome the biological and technical noise in these experiments, researchers have used consensus signatures - averages of multiple experiments - to summarize the effects of perturbations. In this work, we demonstrate that consensus signatures on the L1000 Connectivity Map show a pervasive similarity bias: as more signatures are averaged, the resulting consensus signatures are increasingly similar to each other, regardless of whether the signatures are related. We show that the distribution of Pearson’s correlation changes as a function of the number of signatures averaged. The artifactual similarity bias is caused by skewness in the data and a consequence of using median normalization on non-normal distributions. Furthermore, we show that mean normalization can partly remedy this similarity bias and improve power to identify associations. The similarity bias introduced by consensus signatures is an important potential confounder of analysis of perturbational datasets, and our practical solution could easily be applied by practitioners in the field to improve the analysis of the L1000 Connectivity Map.

## Introduction

Perturbational datasets are a commonly used tool for rapid, high-throughput measurement of the biological changes induced by compounds and genetic reagents [1–3]. Datasets have been generated for several different data modalities and for a number of assays. Characterizing the functional changes induced by perturbations has many applications, like identifying mechanism of action, nominating putative gene agonists and antagonists, constructing biological networks, and identifying differences in samples and cell lines [4–9]. High-throughput perturbational experiments are a critical tool in biomedical research, and any improvement would pay dividends.

Biological perturbational data present analytical challenges because of the low signal-to-noise ratio of the measurements. Biological systems are stable dynamical systems with a preference for homeostasis, and perturbation sufficiently far outside of the stable regime inevitably leads to cell death [10–12]. Additionally, the typical measurements of the biological modalities are high dimensional, as with transcriptomics, proteomics, morphology, and chromatin state, which leads to noise far outweighing the signal.

One commonly-used method to mitigate low signal in perturbational data is to generate consensus signatures: to average multiple independent experiments of the same perturbation under the same or similar conditions [4, 13–23]. The idea is that consensus signatures can leverage the law of large numbers to better estimate the mean changes induced by a particular reagent. In a typical analytical pipeline, similarities are then computed among consensus signatures to identify reagents that produce similar phenotypic changes.

In this paper, we demonstrate an unexpected similarity bias introduced by consensus signatures on the L1000 Connectivity Map that presents a confounder to analyses. We show that this bias emerges from the use of median normalization on skewed data, and we demonstrate that this similarity bias reduces statistical power to detect similar perturbations. This analysis highlights the importance of caution when employing robust normalization on non-normal data and on averaging signatures in high dimensional spaces. To address this issue, we developed a practical correction and use mean normalization to mitigate this phenomenon.

## Results

### Bias in metasignature similarity

For this analysis, we define a metasignature as a feature-wise average of several perturbational experiments. In the literature, this is termed a ‘consensus signature’ [1], though because the definition may vary, we use ‘metasignature.’ The metasignature size (metasize) is the number of component signatures that are averaged. To illustrate the potential pitfalls of analysis with metasignatures, we calculate the similarity of averages of signatures in L1000. We consider the similarity of pairs of disjoint metasignatures of size *k* from three classes: (1) signatures of the same compound treatment, (2) random subsets of all compound signatures, and (3) control signatures of DMSO treatment (Figure 1). Disjoint here means that the component signatures used to average the pair of metasignatures are distinct. The null correlation introduced by averaging is shown by sampling from all compounds and DMSOs - unrelated perturbations and negative controls, respectively - which have no biological association. We compute Pearson’s correlation (PCC) as the similarity metric between pairs of metasignatures for each of the three classes and sample metasignatures 100 times for each value of size to produce a statistical sample.

**Figure 1:**
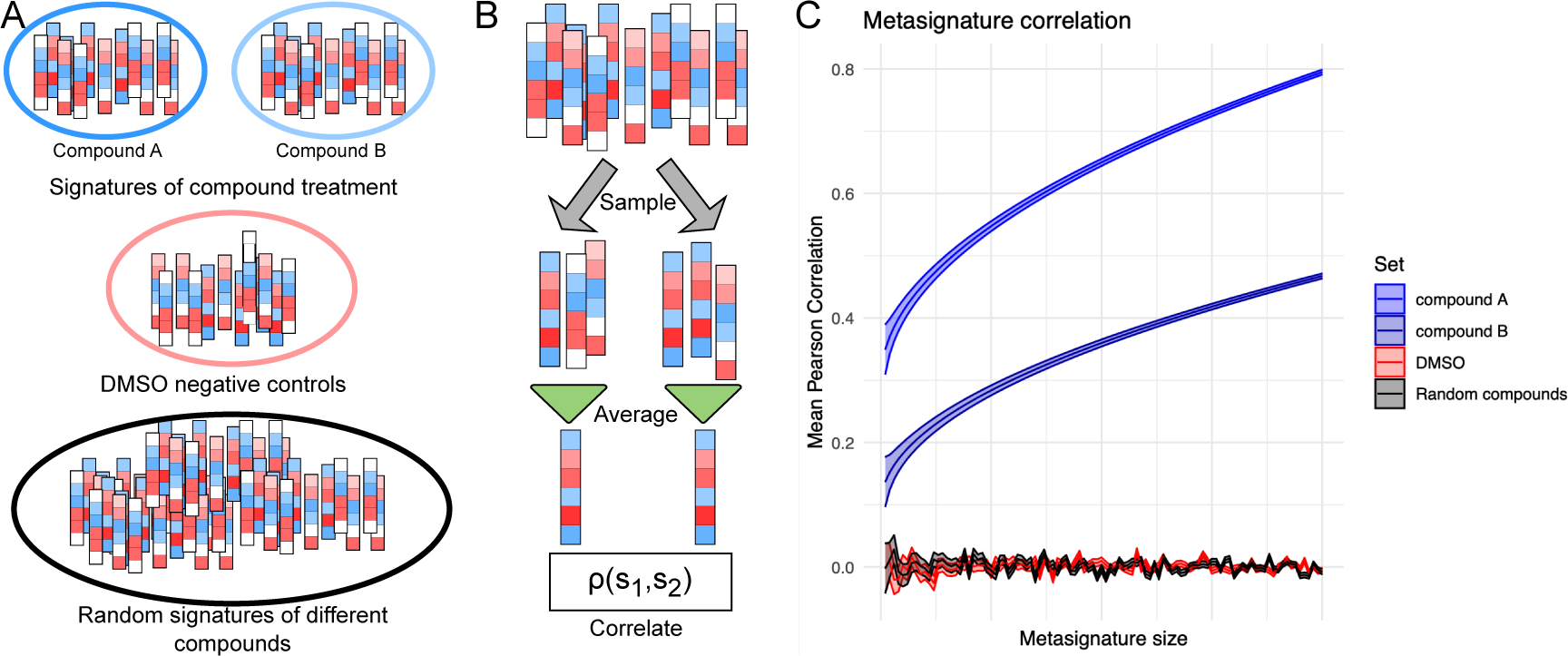
Schematic of the metasignature similarity analysis. Perturbational experiments, like L1000, assay many compounds multiple times - in different cell lines, doses, and time points. (A) We consider sets of signatures for three conditions: a set for each compound treatment, DMSO negative controls, and a random set irrespective of treatment. (B) For a given set, two disjoint subsets of *k* signatures are sampled and averaged, and their Pearson correlation is computed as a function of *k*. (C) The premise of using metasignatures is that the correlation of compound-specific metasignatures will increase as more signatures are averaged because of improved signal-to-noise, whereas the negative controls will not show this increase.

The motivation for averaging is that the metasignatures for a particular chemical compound will better capture the changes induced by that compound. The evidence for this would be better agreement (i.e., higher Pearson’s correlation) among *k* disjoint metasignatures as the size increases. Put another way, averaging more experiments of the same condition should produce a more consistent measurement of the changes induced by that condition. The results show that as the size increases, the Pearson’s correlation for metasignatures of a compound increases (Figures 2a, 2b). However, the null cases of random compounds and DMSOs also show an increase in Pearson’s correlation as a function of the size. This result reveals that there is a systematic bias in similarities between metasignatures that is metasize dependent. Averages of multiple perturbational signatures tend to produce the same result as the number of component signatures increases, whether the component signatures are functionally related or not. Importantly, the distribution of similarities between metasignatures changes as a function of the metasignature size, which presents a major confounder when comparing similarities among metasignatures of varying sizes (Figure 2c). A similarity between two metasignatures of size *N* is necessarily a different distrubition than one between metasignatures of size *M*.

**Figure 2:**
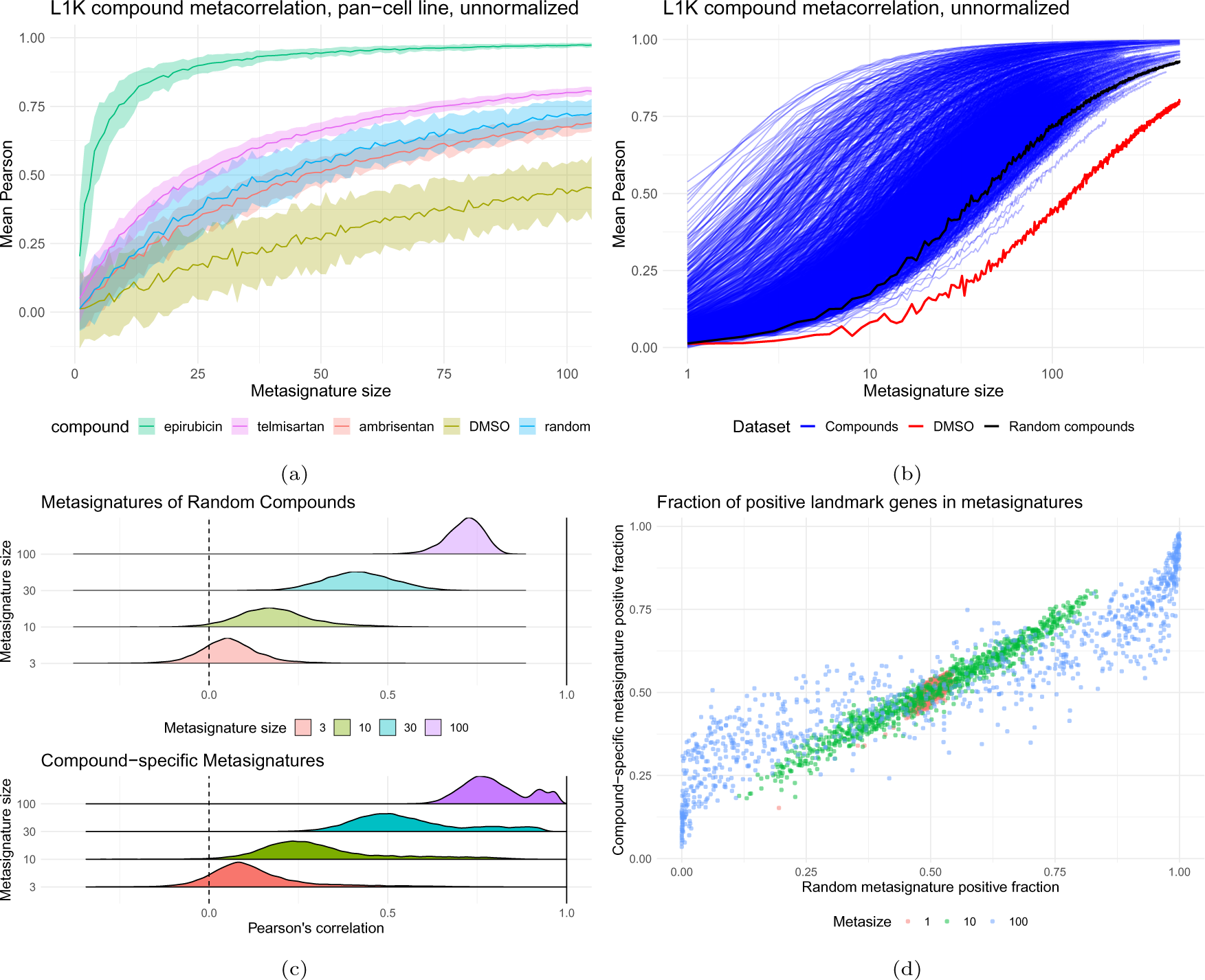
Metasignatures show a positive similarity bias for both compounds and for negative controls as the number of component signatures increases. (a) While compound-specific metasignatures show increasing correlation with size, so do the negative controls DMSO and random compounds. The three example compounds represent the 5th, 50th, and 95th quantiles of compounds by metasignature correlation. (b) The similarity bias is shown for all compounds with at least 100 signatures. Each blue line represents a different compound, and the black and red lines are random compound and DMSO negative controls. Note the log x-axis. (c) The distribution of Pearson’s correlation changes with metasignature size, and increases substantially for random compounds. (d) As more signatures are averaged, the distribution of the signs of the genes switches from approximately 50/50 to more uniformly positive or negative. A gene that is positive in all metasignatures is uninformative.

The reason for the bias in the similarity can be seen by measuring for each feature (the landmark genes in L1000) the fraction of metasignatures with a positive value as a function of size. For differentially expressed genes in a compendium like L1000, the gene is expected to be upregulated about as often as it is downregulated. L1000 uses robust normalization, where the data is centered on the median and scaled by the median absolute deviation. Level 4 data from L1000 by construction has median 0, so the fraction of Level 4 instances with a positive value for each gene is 0.5. We considered the fraction of positive values for each gene for (1) compound-specific metasignatures and (2) random metasignatures, with metasignature sizes of 1 (i.e. unaveraged), 10, and 100. As the metasignature size increases, the positive fraction tends to 0 or 1, and critically, the tendency is the same for both compound-specific and random metasignatures (Figure 2d). For size of 1, the genes are positive close to 50% of the time; for metasignatures of size 100, 330/978 genes are positive or negative at least 80% of the time for compounds, and 927/978 genes are positive or negative at least 80% of the time for random sets. Moreover, the consistency of the sign bias in gene features indicates that both compound-specific and random metasignatures tend towards the same value.

### Causes of the bias

The change in the similarity distribution suggested that averaging was affecting the underlying gene-level distributions. To investigate this, we considered the moments of each measured gene over the space of signatures for metasignatures of size 1, 10, and 100. Unsurprisingly, the distribution of the gene-wise mean is unchanged by averaging, and the standard deviation of the genes goes as roughly *n^−^*^1*/*2^, where n is the metasignature size (Figure 3a). What is surprising is that the distribution of Mean z-score is dramatically non-zero for most of the genes, when the genes were normalized with robust normalization. This observation also holds within each cell line (Figure S1a).

**Figure 3:**
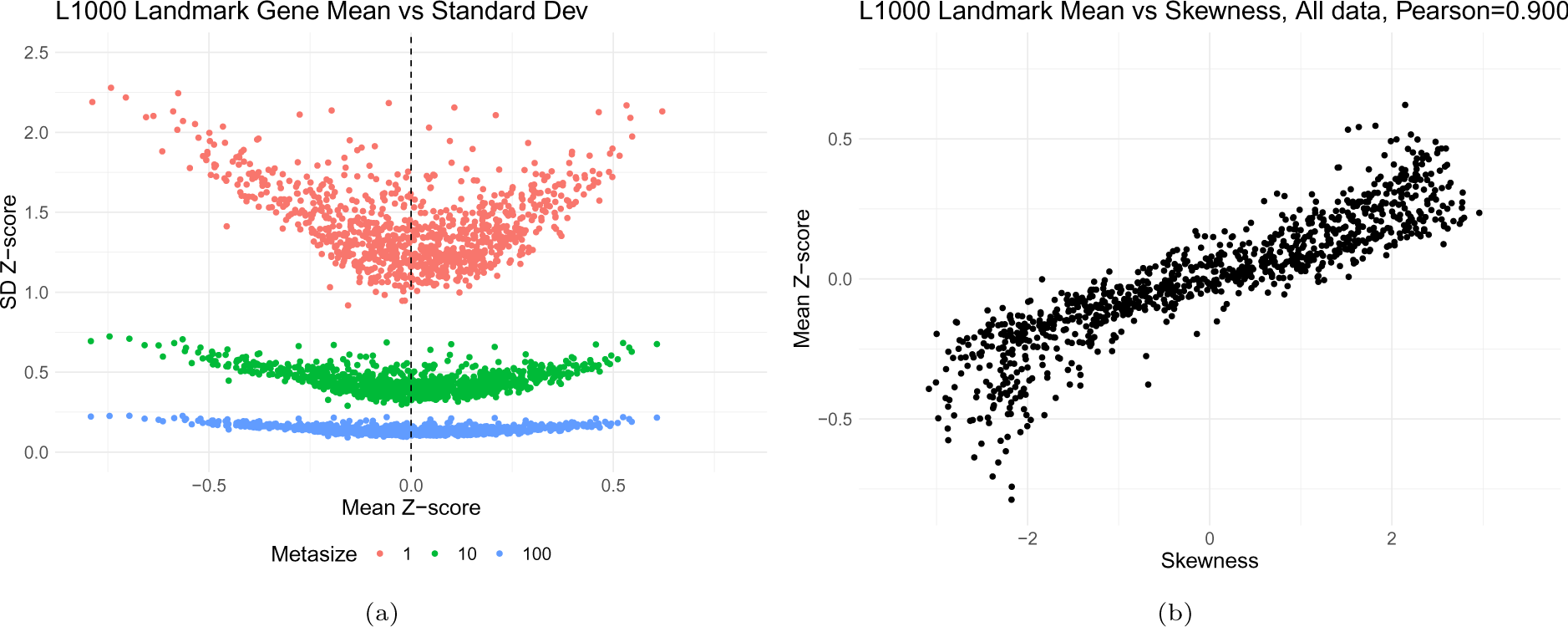
The moments of L1000 Level 5 dataset for each of the landmark genes. (a) Shows the mean Z-score vs the standard deviation Z-score for each landmark gene over the L1000 compound signatures and random metasignatures. By construction in L1000, a Z-score of 0 is presumed to indicate no modulation by a perturbation, but many of the landmark genes have mean Z-scores with absolute magnitude greater than 0.5. As signatures are averaged into metasignatures, the mean stays constant while the variance decreases. (b) The skewness of the landmark genes over compound data vs the mean Z-score. Median centering of skewed data results in non-zero mean, which is the root cause of the observed similarity bias.

We reasoned then that while the median gene expression was set to 0, the distributions had non-zero mean. Robust normalization tacitly assumes a roughly normal distribution with a few outliers, and median centering is motivated by robustness to these outliers. However, the distributions of L1000 genes deviate substantially from normality, and in particular have non-zero skew (Figure 3b). The tendency of skew distributions is for the mean to be greater than the median for positive skew and less than the median for negative skew. We found that the skewness of measured landmark genes for unaveraged level 5 signatures correlated very well with the mean, with a Pearson’s correlation of 0.9007. The observation also holds in cell line-specific data, where the Pearson’s correlation is 0.8391 (Figure S1b).

Furthermore, the skewness of the data is not simply caused by the outliers, as is a premise of robust normalization. To illustrate this, for all L1000 compound signatures in landmark gene space, we computed the skewness of each of the 978 landmark genes and compared this against the skewness of the inner 90% of the data - the data between the 5th and 95th quantile - excluding the most extreme 5% of the data at each tail. If the skewness were driven purely by these most extreme 10% of the data points with an otherwise normal distribution, the 90th percentile skewness would be 0. We find that while the skewness of the inner 90% of the data is lower (as expected when removing outliers), it is non-zero (Figure S2). The correlation between landmark gene skewness and skewness of the central 90% is Pearson = 0.874.

### Correcting the bias

The working hypothesis is then that the similarity bias is introduced by non-zero mean for genes over the space of signatures, which is in turn a result of median normalization on non-normal distributions. To test this and correct for the similarity bias, we centered the data so that the mean is 0, then evaluated the similarity bias. However, it is unclear whether to use mean of the negative controls (DMSO signatures) or the background of all compounds for centering. We therefore normalized two different versions of the data - for compound normalization, the mean of each landmark gene over the space of compound signatures was subtracted and set to 0; for DMSO normalization, the mean for DMSO signatures was set to 0. We then evaluated the similarity bias for all three cases as before, but with the renormalized data. We found that the similarity bias for the negative control use for centering vanished, i.e. metasignatures from that control showed no increase in Pearson’s correlation, suggesting that the offset mean hypothesis as a driver of similarity bias is correct (Figure 4a, 4b).

**Figure 4:**
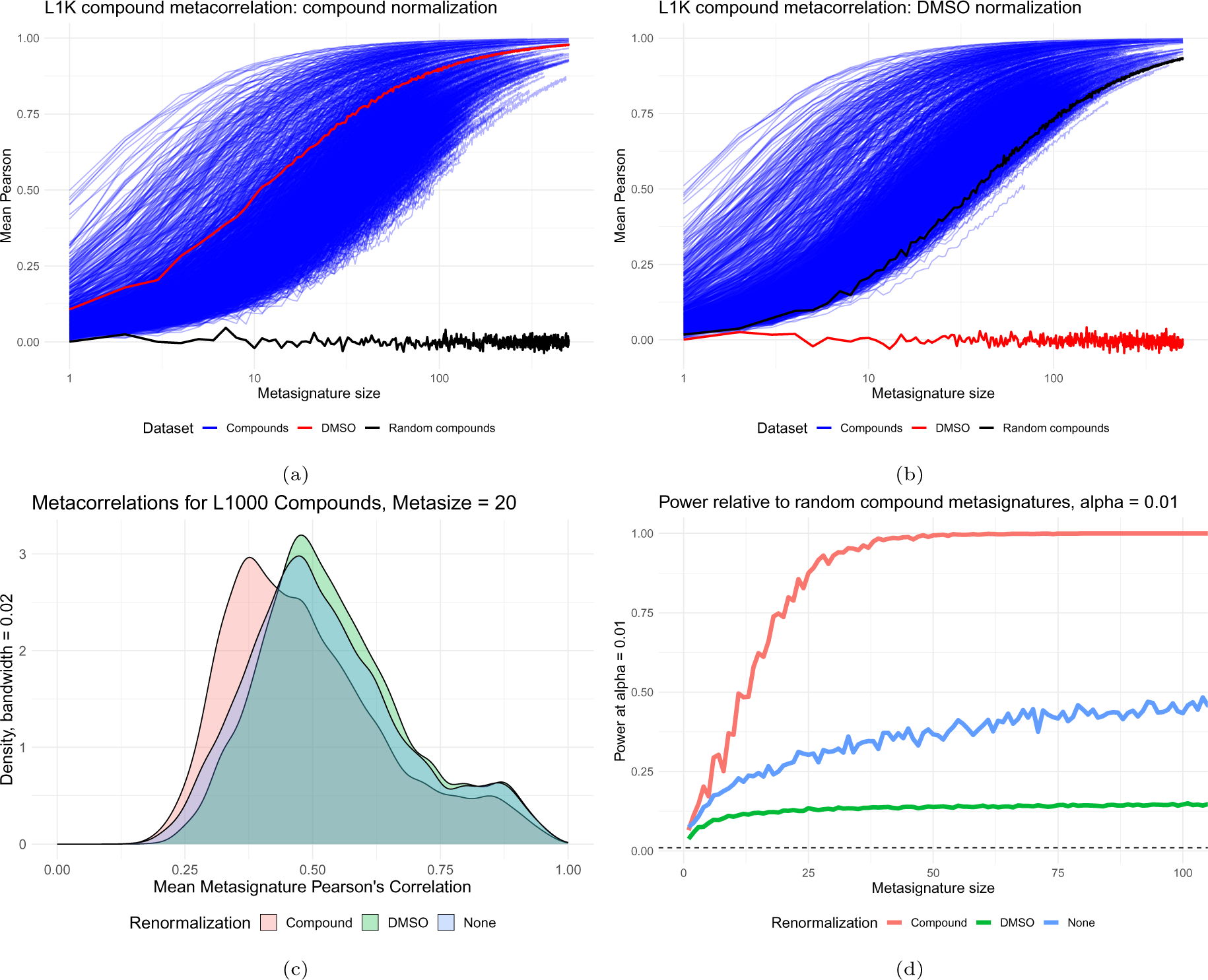
Renormalizing the data relative to the negative controls removes the similarity bias in those controls. (a) Recentering the data so that the mean Z-score over the space of all compounds is zero for each gene removes the similarity bias for metasignatures from random compounds. (b) Recentering the data on the DMSO signatures removes the similarity bias for DMSO metasignatures. The two negative controls do not simultaneously correct each other. (c) The distribution of metasignature correlation changes with renormalization, with a marked decrease with compound normalization, here shown for metasignature size = 20. (d) The statistical power, or fraction of compound-specific metasignatures with correlation greater than (1-*α*) of random metasignatures, varies for the renormalizations.

Surprisingly, when one negative control - random compounds or DMSOs - was centered, the other control showed worse metasignature similarity bias, suggesting that the two nulls do not have the same underlying distribution (Figure S3). We posit that the distributions of the two negative controls are different, whether from a confounder like plate effects [24, 25], a bias in the compounds assayed towards a particular biological effect, or a technical artifact. To choose a normalization scheme, we considered the distribution of metasignature similarities with the presumption that the normalization that reduces the compound-specific metacorrelation the most is more correctly normalizing the data. Compound renormalization most reduces the compound-specific metacorrelation, though the correction is marginal (Figure 4c). Comparison of compound-specific metasignatures to random compounds is also conceptually the most sensible negative control, presuming that the distribution of assayed compounds is sufficiently diverse.

While it is possible to control the similarity bias for negative controls through renormalization, the tendency for compound-specific metasignatures to show consistently increasing correlation is an unexpected result. While there are strong bioactives in the L1000 dataset, at least some fraction of the compound treatments are expected to be inert, for instance some from the Diversity Oriented Synthesis library[26]. To quantify the fraction of compounds with a measurable signal, we compared compound-specific metasignature correlation to that of random compounds for each normalization and measured what proportion of metasignature correlations were detected at *α* = 0.01. That is, we measured how often compound-specific metasignatures have a correlation greater than the top 1% of null correlations. Based on the assumption that not all compounds are bioactive, we expect that the correct recentering will recover some large fraction of compounds, but not all. At a metasignature size of 50, DMSO renormalization has power of 0.138, the base dataset has power of 0.367, and compound renormalization has power of 0.994 (Figure 4d). This result suggests that with mean normalization of the set of all compound signatures, a metasignature of 50 averaged experiments for a compound is sufficient to identify that compound from the database. Most compound perturbations will not have 50 signatures in any perturbational experiment. At metasignature size of 10, DMSO normalization has a power of 0.108, base dataset has 0.209, and compound normalization has 0.366 (Table S1). This shows that using mean normalization approximately doubles the power to identify compound treatments based on the signature compared to the median normalized dataset.

Because the compound renormalization forces the compound mean to 0 before using random draws as negative controls, we posited that the result was circular. To test this, we separated the normalization and negative control step. First, we recentered the dataset using the mean expression in half of the compound signatures. We then used the other half of the compound signatures to evaluate the correlation among random metasignatures. We found that the similarity bias was greatly reduced in the excluded portion of the dataset not used in recentering, suggesting that the recentering is not circular (Figure S4).

### Offset power analysis

While the biases from metasignatures are apparent, it is not obvious that this confounds analysis - the goal of which is to identify signatures of reagents that produce similar transcriptomic changes. To explore the effect of a non-zero mean on power, we artificially added random offsets to signatures from the ten most assayed cell lines in L1000, using the basic level 5 perturbational signatures rather than averaging to create metasignatures. We computed the power to discriminate correlations between signatures of the same compound from similarities between all pairs of signatures. We then observed the change in power as the data was increasingly offset from 0 mean. Our hypothesis was that the power, recall, and z-scores would be maximal when the mean was 0, and would deteriorate as the offset was increased.

We observed that statistical measures tended to decrease as the offset increased, but were not always maximized when the offset was 0. The power - the fraction of same-compound correlations ranking in the top 1% relative to the background - was greater for several cell lines with offsets of 5-10 standard deviations before declining as the offset grew sufficiently large (Figure 5a). Recall at a particular FDR showed a similar trend, peaking around 5-10 standard deviations for several cell lines (Figure 5b, 5c). The fraction of compound pairs with Z-scores *>* 2 consistently decreased with increasing offset (Figure 5d). This analysis was a simple illustration rather than an exhaustive exploration of the impact of offsets in the data, but it suggests that simply mean centering the data, although highly beneficial, may not maximize analytical power of perturbational datasets like L1000.

**Figure 5:**
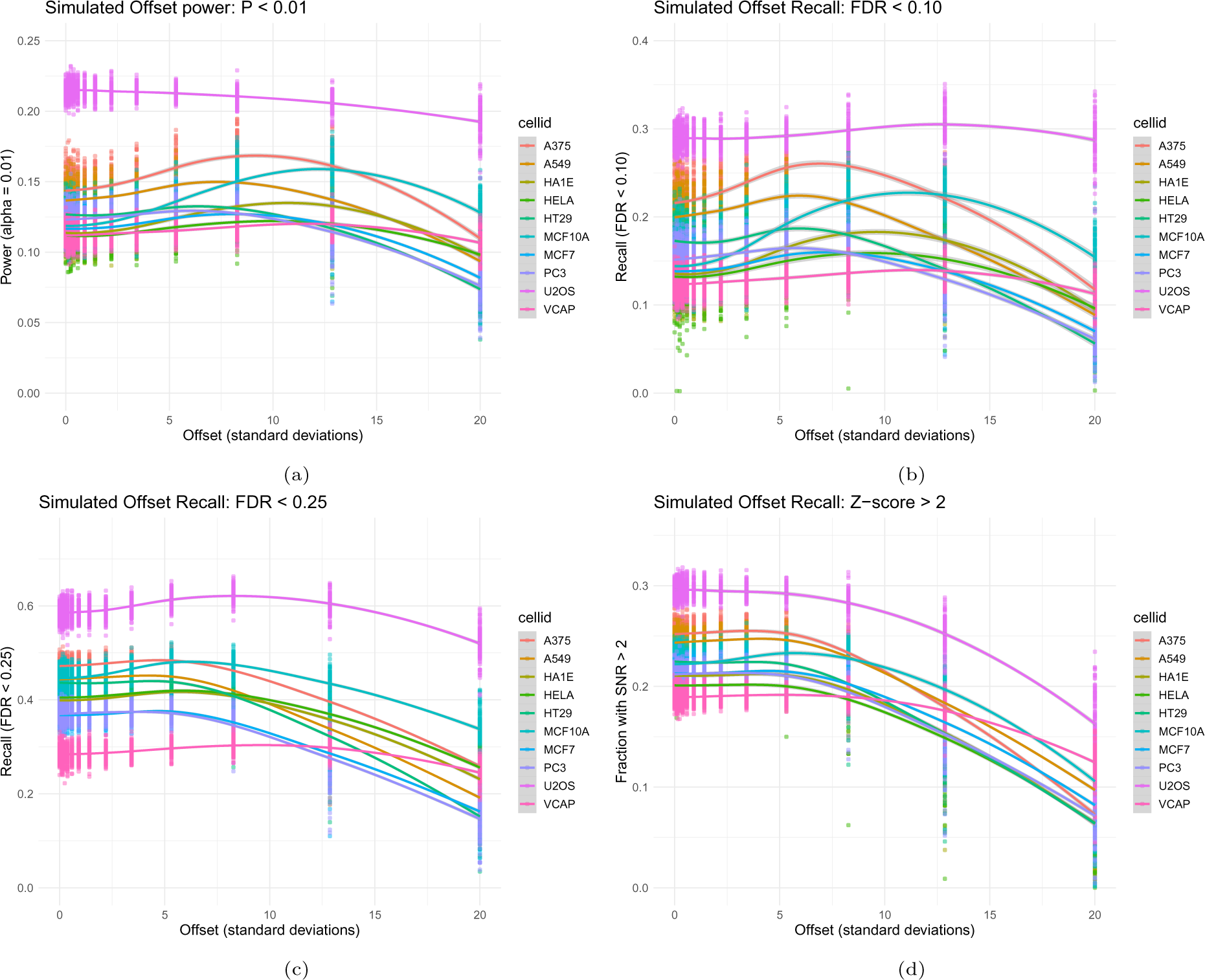
Introducing shifts to the mean of the distribution affects the statistical power and recall of similarities between compound signatures in the L1000 dataset. The Pearson’s correlation is computed among signatures of the same compound, and this is compared to a null distribution of all signatures. We then compute the (a) power at *α* = 0.01, (b) recall at FDR *<* 0.1, (c) recall at FDR *<* 0.25, and (d) fraction of positive pairs with Z-score *>* 2.

## Discussion

In this work, we have identified a bias where the correlation between metasignatures (or consensus signatures) on the L1000 dataset of increasing size increases systematically regardless of the composition of the metasignature - whether compound specific, permutation null, or negative control. The distribution of similarities - correlation, cosine similarity, or gene-set methods - changes with the number of component signatures. This similarity bias is a significant confounder when comparing similarities among perturbational signatures, especially when naively ranking similarities relative to each other. Averaging more signatures together converges to a common average, which limits power to discriminate signatures and changes the distribution of similarities.

Furthermore, we identified the underlying cause of this detrimental phenomenon: the mean of the features (gene expression) is not zero. The L1000 dataset uses a median normalization that sets the median to zero. However, because many of the genes have skewed, non-normal distributions, it leads to non-zero mean expression for many genes or features. Averaging signatures shrinks the result toward this mean, leading to different similarity distributions with consensus signatures. This similarity bias suggests that median normalization can introduce artifactual offsets in a dataset because the implicit assumption of normality breaks down. Naive commingling of similarities among consensus signatures of different sizes will result in an increase in both false positives and false negatives because the similarity distributions are different. Our suggestion is to compare any similarities computed between metasignatures against a null distribution of the same size metasignatures.

We proposed and evaluated a corrective solution, i.e. normalizing the data so the mean expression for compound signatures is zero. A key result is that mean centering substantially improves the power to match two metasignatures of the same compound treatment compared to median centering. The discriminative power of compound-specific metasignatures of size 100 increases to 1 with compound normalization, which is a result unlikely to have biological basis, as not all compounds are expected to be bioactive. However, in practice, the vast majority of compound consensus signatures will be of much smaller size because they have fewer repeated experiments.

In summary, particularly in noisy, high-dimensional biological spaces, averaging perturbational signatures is a seemingly sensible strategy to identify changes induced by perturbation that can lead to negative consequences that drastically reduce the utility of these experiments. Caution must be used when employing consensus signatures to ensure the statistics are properly treated and the normalization method is suitable.

## Methods

### Data

We used the L1000 dataset originally published in [1]. We used the level 5 dataset of compound and vehicle (DMSO) signatures from the 2020 release accessed via the Broad Institutes’s web portal at http://clue.io. We considered only the 978 directly measured landmark transcripts, ignoring the computationally inferred genes. The data was manipulated with the cmapR package, and additional functionality is found in the BHK Lab’s perturbKit package.

The median normalization procedure, as described in [1], is as follows, which we reproduce here for convenience. The key point is that the *median* of a gene is centered, which for skew data typically leads to a non-zero mean.

To obtain a measure of relative gene expression, we use a robust z-scoring procedure to generate differential expression values from normalized profiles. We compute the differential expression of gene *x* in the *i*th sample on the plate as:

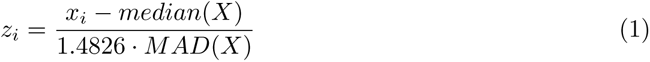

where X is the vector of normalized gene expression of gene x across all samples on the plate, MAD is the median absolute deviation of X, and the factor of 1.4826 makes the denominator a consistent estimator of scale for normally distributed data.

### Bias in metasignature similarity

Exploring the changes in similarity for metasignatures requires generating large numbers of metasignatures. To do this, for each compound, we selected all signatures with that compound treatment irrespective of dose, cell line, and time point. We then sampled k of these without replacement and averaged them to generate a metasignature of size *k*. To compute the similarity bias, we sampled two disjoint metasignatures of the same size - disjoint meaning they have no common component signatures. We then used Pearson’s correlation over the 978 landmark genes. For comparison, we also evaluated metasignature correlation using the DMSOs and using random draws from all compounds.

The fraction of positive landmark genes in metasignatures quantifies for each landmark gene what fraction of metasignatures have a positive value for that landmark. For example, for some gene G, and some metasignature size *k*, we constructed metasignatures of size *k*, then calculated what fraction of those metasignatures had a positive value for G. For the random metasignatures, we sampled 100,000 random signatures, then exhaustively constructed disjoint signatures of size 1, 10, and 100. For the compound-specific metasignatures, we used the 740 compound treatments with more than 200 signatures in the dataset, and exhaustively constructed disjoint signatures of size 1, 10, and 100. The motivation of this analysis was to characterize gene-specific biases induced by metasignatures. Any deviation from roughly 50% positive values indicates that metasignatures are systematically biased for that gene. That the compound-specific metasignatures and random metasignatures show the same bias for each gene indicates that this bias is an artifact and not specific to the compound treatment.

### Moments of the dataset

The investigation into the causes of the similarity bias required computing moments of the landmark genes from the L1000 Level 5 signature data. We used base R and the moments package to compute the mean, variance, and skewness of each gene across the space of signatures or metasignatures.

### Corrective normalization

The purpose of renormalizing the data is to remove the non-zero mean introduced by applying median normalization to non-normal data. The base data, originally published in [1] is referred to as “None” renormalization in this manuscript. The two normalization schemes we applied are compound centering and DMSO centering. For compound centering, we subtracted the mean over all compound signatures from each gene, so that the resulting data set had gene scores with zero mean. When analyzing the entire dataset (Figure 4a), we normalized with all the signatures, though in principle, it is reasonable to analyze each cell line independently. For DMSO centering, we computed the mean of each gene across the DMSO signatures and subtracted this from the total dataset, meaning that the mean expression on the renormalized DMSO signatures was zero.

### Metasignature power

The statistical power of a dataset and similarity metric (Pearson, throughout this paper) is the probability of rejecting the null hypothesis when the alternate hypothesis is true. In this case, we assume that the alternative hypothesis is true for two disjoint metasignatures of the same compound - that is, that they represent the same biological effect. The null distribution by assumption is the distribution of Pearson’s correlations between random metasignatures, i.e., averages of collections of unrelated compound signatures randomly sampled from the dataset. We reject the null hypothesis at significance level *α* if the correlation between two metasignatures of the same compound exceeds all but *α* of the null hypothesis. So, for *α* = 0.01, the dataset and similarity metric are powered to detect the fraction of same-compound metasignatures with correlation in the top 99% of the null distribution. For the power analysis, we considered the 752 of 33609 compounds (using the “pert iname” identifier from the L1000 metadata) with at least 200 signatures. This allowed a power analysis that considered the same set of compounds over the range of metasizes 1-100.

### Offset power

The offset power analysis was a simulation to see how introducing a non-zero mean along various axes affects discriminative power. This analysis did not use metasignatures, but just computed Pearson’s correlation among level 5 signatures in the landmark gene space. Technically, these are equivalent to metasignatures of size 1. First, we sampled a random direction in landmark space and set the mean of the data along that axis to zero. We then shifted the data along that axis by twelve different values in units of the standard deviation of the data along that axis. We computed the power to identify signatures of the same compound compared to the background similarity, recall at FDR = 0.1 and 0.25, and the Z-score of Pearson’s correlation of signatures of the same compound relative to the background. We repeated the process 100 times with different directions. We applied this analysis to the ten cell lines in the L1000 data with the most compound signatures.

## Key Points

- Averaging perturbational signatures from L1000 changes the distribution of similarities between signatures and introduces a positive similarity bias that persists for unrelated consensus signatures.
- The similarity bias of consensus signatures depends on the number of component signatures and is a consequence of using median normalization on skewed, non-normal distributions for gene expression.
- Mean centering the data can mitigate the similarity bias and substantially improve power to detect related signatures.
- Use of consensus signatures with median normalization and comparing similarities of consensus signatures irrespective of the number of component signatures can increase errors and reduce power on perturbational datasets.

## Glossary

**metacorrelation** The correlation (Pearson’s) between pairs of metasignatures. 6

**metasignature** The average of k signatures, also called “consensus signature”. 2

**metasize** The number of component signatures in a metasignature. 2

## Author Contributions

I.S. and B.H.K. conceived of and planned the study. I.S. designed and executed the analysis. I.S. wrote the manuscript with contributions from K.S. and B.H.K. I.S. and K.S. developed the software. B.H.K. supervised the study and refined the analysis.

## Acknowledgments

We thank the Broad Institute and the LINCS Consortium for sharing the L1000 data with the scientific community.

## Software

The analysis was run in R 4.2.1 on a high performance computing server. We used the following R packages, available from Bioconductor unless otherwise noted.

- cmapR 1.10.0 - CMap Tools in R, [27] I/O and manipulation of L1000 datasets
- metasignatures 0.900 - metasignatures, a BHK Lab package for manipulation of metasignatures, available at https://github.com/iamsinht/metasignatures
- perturbKit 0.900 - perturbKit, a BHK Lab package for manipulation of perturbational datasets, available at https://github.com/bhklab/perturbKit
- ggplot2 3.4.3, MASS 7.3.60, moments 0.14.1, stats 4.2.1, coop 0.6.3

## Data Availability

The analysis data underlying this article are available in the article and in its online supplementary material. The data for the L1000 dataset are in the public domain and are available from https://clue.io/[1].

## Supplemental Figures

**Figure S1:**
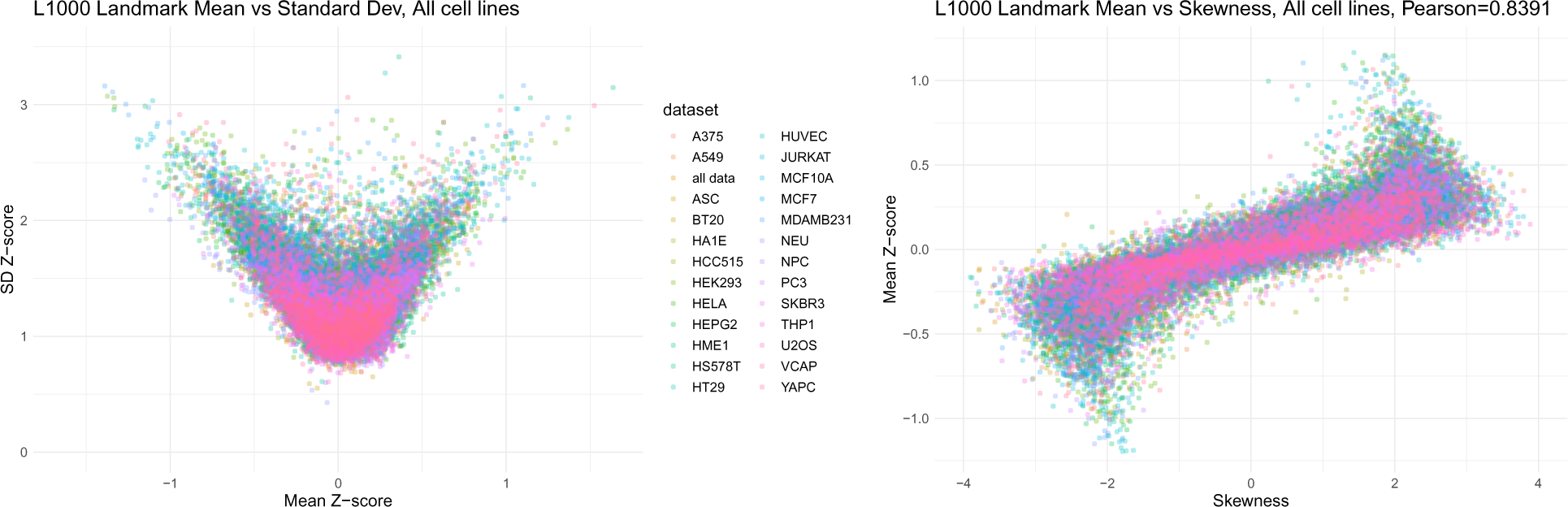
Moments for L1000 signatures for a set of the most assayed cell lines. Each point represents a landmark gene in one of the cell contexts. (a) The mean and standard deviation of the Z-scores for compound signatures show that many landmark genes have non-zero mean value despite the robust normalization. (b) The relationship between mean and skewness for each cell line shows a strong correlation, with Pearson = 0.8391.

**Figure S2:**
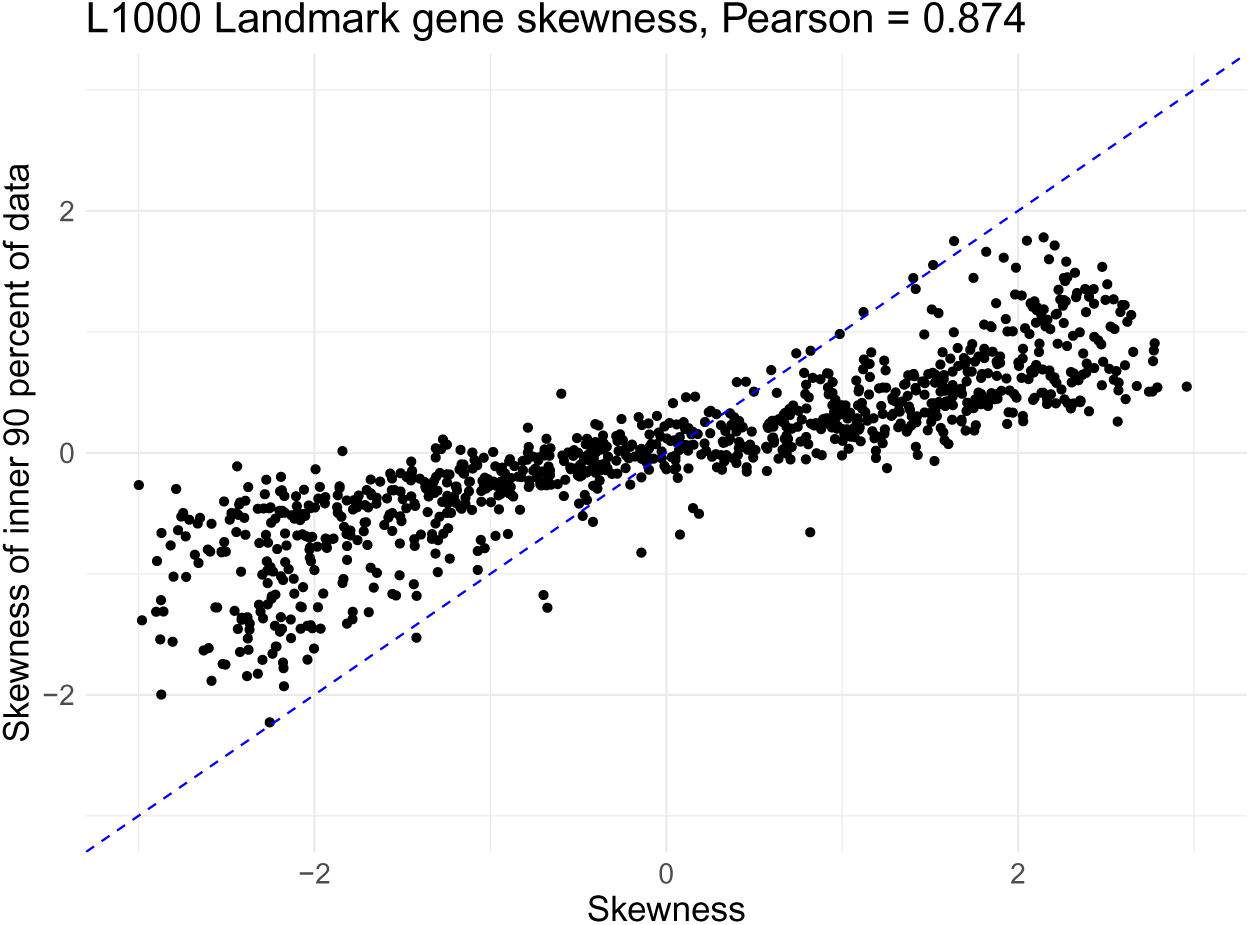
For each landmark genes, for all level 5 compounds, this figure shows skewness of each gene plotted against the skewness of the data in the inner 90% of the data - between the 5th and 95th quantiles. An assumption of robust normalization is that the core of the data is normally distributed with a small number of outliers. This shows that while the most extreme 10% of the data does contribute to the skewness, the inner 90% of the data still has non-zero skewness. This illustrates a source of a pitfall of applying median-centered robust normalization to non-normal data.

**Figure S3:**
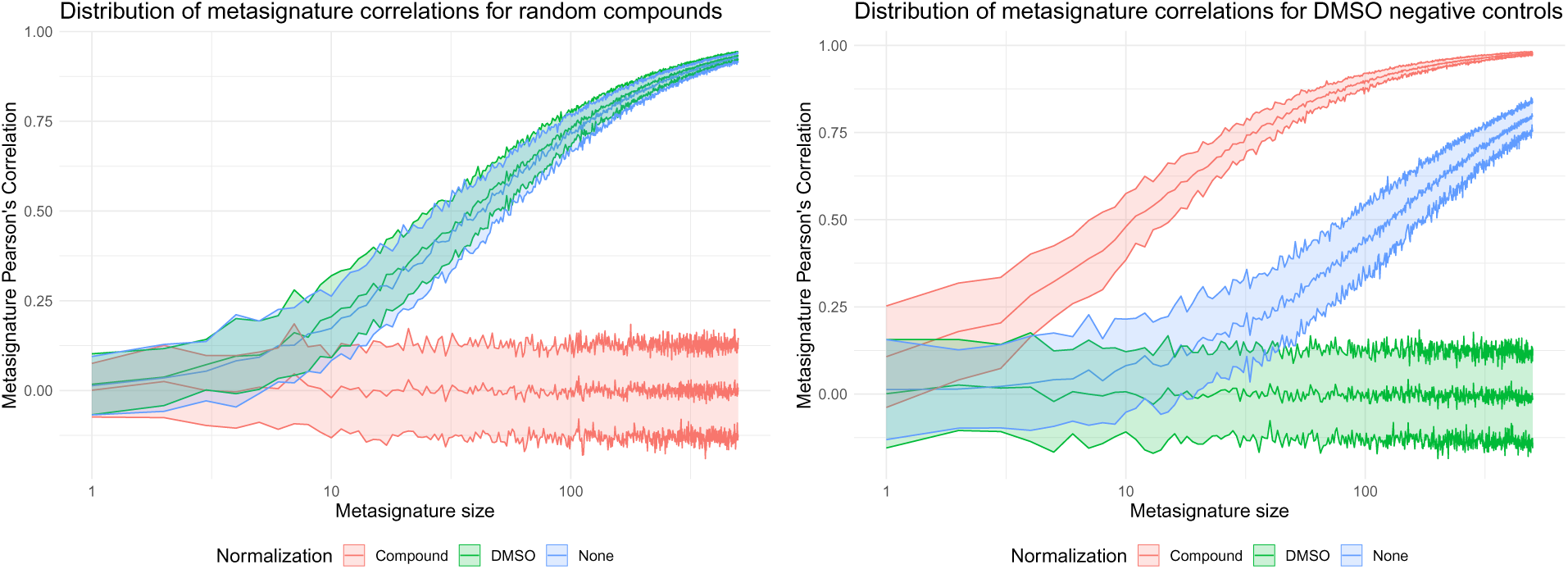
Renormalizing with respect to all compounds or to DMSO signatures changes the metasignature correlation curves for random compounds (left) and DMSOs (right), both of which are negative controls. Curiously, normalizing by one increases the similarity bias of the other, suggesting the distributions for the two negative controls are not the same.

**Table S1:**
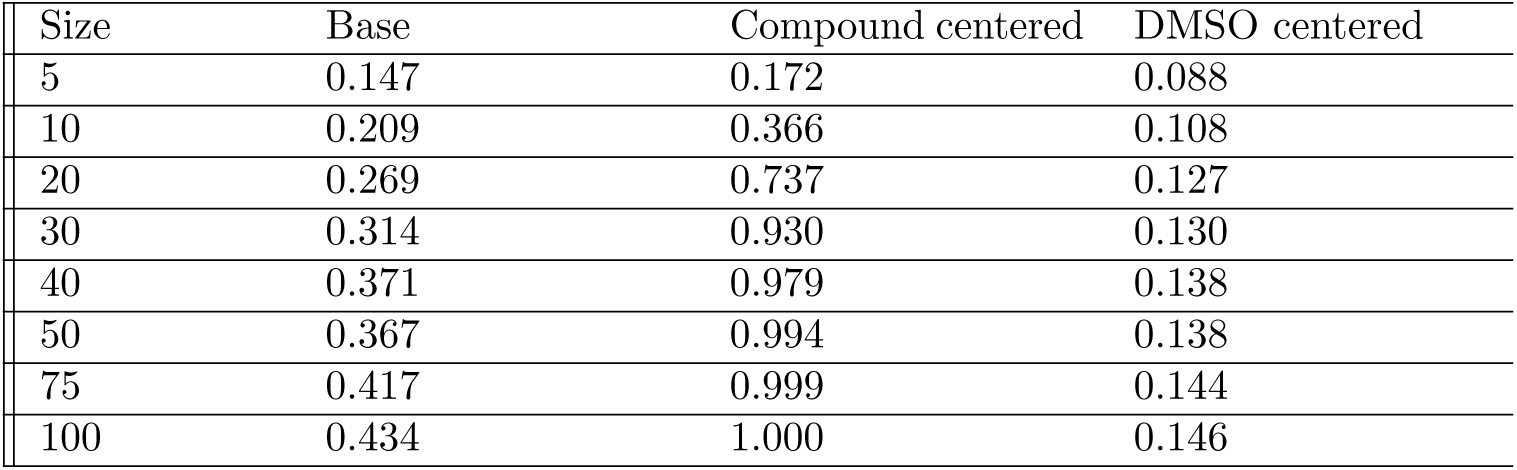
Table of power to connect two disjoint metasignatures of the same compound at *alpha* = 0.01 for different normalization schemes. These data correspond to Figure 4d. For each value of the size of metasignatures, the power is the fraction of disjoint compound-specific metasignatures with Pearson’s correlation greater than (1 *− α*) of random metasignatures.

**Figure S4:**
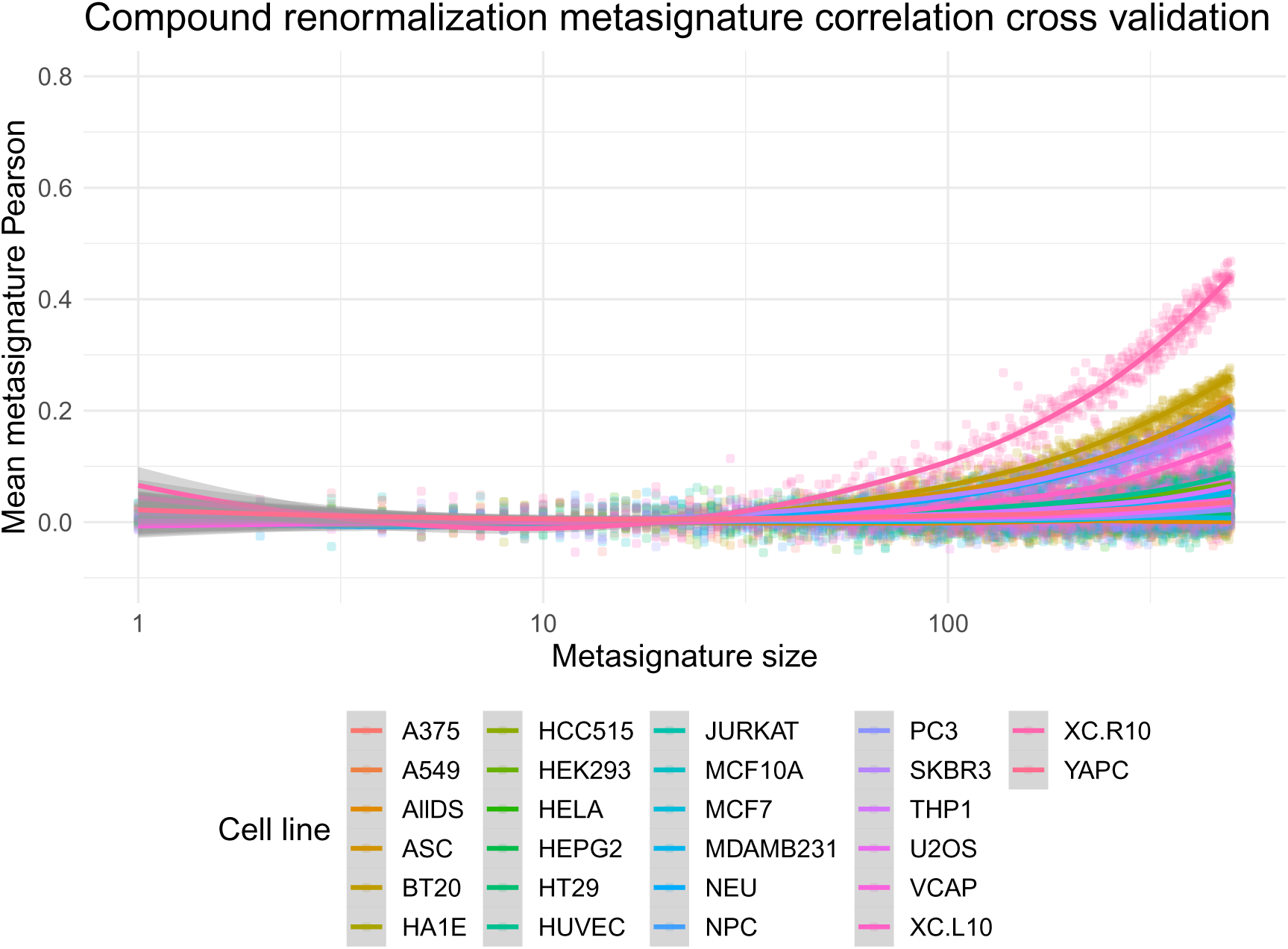
Compound recentering on a separate dataset still results in an overall decline in random compound metasignature similarities. Here, we used half the data to recenter, so the mean was zero on that set of compound signatures. We then used the other half, random generated metasignatures, and computed the Pearson’s correlation with increasing metasignature size. Even with separate renormalizing and metasignature similarity sets, the similarity bias is greatly reduced (contrast with Figure 2b). Only XC.R10, BT20, SKBR3, and ASC have metasignature correlation *>* 0.1 for metasize = 200, perhaps indicating data heterogeneity in those cell lines.

## References

1. Subramanian, A. et al. A Next Generation Connectivity Map: L1000 Platform and the First 1,000,000 Profiles. Cell 171. arXiv: http://biorxiv.org/content/early/2017/05/10/136168 Publisher: Elsevier, 1437–1452.e17. issn: 00928674. http://www.ncbi.nlm.nih.gov/pubmed/29195078 (2017) (Nov. 2017).

2. Lamb, J., et al. The connectivity map: Using gene-expression signatures to connect small molecules, genes, and disease. Science 313. Publisher: American Association for the Advancement of Science, 1929–1935. issn: 00368075. (2020) (Sept. 2006).

3. Bray, N. L., Pimentel, H., Melsted, P. & Pachter, L. Near-optimal probabilistic RNA-seq quantification. Nature Biotechnology 34, 525–527. issn: 15461696. http://www.nature.com/ (2019) (2016).

4. Niepel, M. et al. Common and cell-type specific responses to anti-cancer drugs revealed by high throughput transcript profiling. Nature Communications 8. Publisher: Nature Publishing Group, 1186. issn: 20411723. http://www.nature.com/articles/s41467-017-01383-w (2017) (Oct. 2017).

5. Corsello, S. M. et al. Bryan 1,6, Ranad Humeidi 1, David Peck 1. bioRxiv. Publisher: Cold Spring Harbor Laboratory, 730119. https://www.biorxiv.org/content/10.1101/730119v1 (2019) (Aug. 2019).

6. Caicedo, J. C., et al. Data-analysis strategies for image-based cell profiling. Nature Methods 14. Publisher: Nature Research, 849–863. issn: 15487105. http://www.nature.com/doifinder/10.1038/nmeth.4397 (2017) (Aug. 2017).

7. Chandrasekaran, S. N. et al. JUMP Cell Painting dataset: morphological impact of 136,000 chemical and genetic perturbations en. Pages: 2023.03.23.534023 Section: New Results. Mar. 2023. https://www.biorxiv.org/content/10.1101/2023.03.23.534023v1 (2023).

8. Way, G. P. et al. Morphology and gene expression profiling provide complementary information for mapping cell state. en. Cell Systems 13, 911–923.e9. issn: 2405-4712. https://www.sciencedirect.com/science/article/pii/S2405471222004021 (2022) (Nov. 2022).

9. El-Hachem, N., et al. Integrative cancer pharmacogenomics to infer large-scale drug taxonomy. Cancer Research 77. Publisher: American Association for Cancer Research, 3057–3069. issn: 15387445. https://pubmed.ncbi.nlm.nih.gov/28314784/ (2019) (June 2017).

10. Vanhaelen, Q., Aliper, A. M. & Zhavoronkov, A. A comparative review of computational methods for pathway perturbation analysis: dynamical and topological perspectives. en. Molecular BioSystems 13. Publisher: The Royal Society of Chemistry, 1692–1704. issn: 1742–2051. https://pubs.rsc.org/en/content/articlelanding/2017/mb/c7mb00170c (2023) (Aug. 2017).

11. Nijhout, H. F., Best, J. A. & Reed, M. C. Systems biology of robustness and homeostatic mechanisms. eng. Wiley Interdisciplinary Reviews. Systems Biology and Medicine 11, e1440. issn: 1939-005X (May 2019).

12. Tyson, J. J. & Novak, B. A Dynamical Paradigm for Molecular Cell Biology. Trends in cell biology 30, 504–515. issn: 0962-8924. https://www.ncbi.nlm.nih.gov/pmc/articles/PMC8665686/ (2023) (July 2020).

13. Schubert, M. et al. Perturbation-response genes reveal signaling footprints in cancer gene expression. en. Nature Communications 9. Number: 1 Publisher: Nature Publishing Group, 20. issn: 2041-1723. https://www.nature.com/articles/s41467-017-02391-6 (2022) (Jan. 2018).

14. Ler Chow, Y. I., Singh, S. I., Carpenter ID, A. E. & Way ID, G. P. Predicting drug polypharmacology from cell morphology readouts using variational autoencoder latent space arithmetic. PLOS Computational Biology 18 (ed Haugh, J. M.) Publisher: Public Library of Science ISBN: 1111111111, e1009888. issn: 1553-7358. https://journals.plos.org/ploscompbiol/article?id=10.1371/journal.pcbi.1009888 (2022) (Feb. 2022).

15. Krug, K. et al. A Curated Resource for Phosphosite-specific Signature Analysis. Molecular and Cellular Proteomics 18. Publisher: American Society for Biochemistry and Molecular Biology Inc., 576–593. issn: 15359484. http://www.mcponline.org/article/S1535947620318600/fulltext (2022) (Mar. 2019).

16. Douglass, E. F., et al. A community challenge for a pancancer drug mechanism of action inference from perturbational profile data. Cell Reports Medicine 3. Publisher: Cell Press, 100492. issn: 26663791. (2022) (Jan. 2022).

17. Way, G. P. et al. Predicting cell health phenotypes using image-based morphology profiling. Molecular Biology of the Cell 32. Publisher: American Society for Cell Biology, 995–1005. issn: 19394586. https://www.molbiolcell.org/doi/abs/10.1091/mbc.E20-12-0784 (2021) (Apr. 2021).

18. Abelin, J. G. et al. Reduced-representation phosphosignatures measured by quantitative targeted MS capture cellular states and enable large-scale comparison of drug-induced phenotypes. Molecular and Cellular Proteomics 15. Publisher: American Society for Biochemistry and Molecular Biology Inc., 1622–1641. issn: 15359484. /pmc/articles/PMC4858944/?report=abstract (2020) (May 2016).

19. Szalai, B. et al. Signatures of cell death and proliferation in perturbation transcriptomics data - from confounding factor to effective prediction. Nucleic Acids Research 47, 10010–10026. issn: 13624962. https://portals.broadinstitute. (2020) (2019).

20. Lin, K., et al. A comprehensive evaluation of connectivity methods for L1000 data. Briefings in Bioinformatics. issn: 1467-5463. https://academic.oup.com/bib/advance-article-abstract/doi/10.1093/bib/bbz129/5626334 (2020) (2019).

21. Méndez-Lucio, O., Baillif, B., Clevert, D. A., Rouquié, D. & Wichard, J. De novo generation of hit-like molecules from gene expression signatures using artificial intelligence. Nature Communications 11. Publisher: Nature Research, 1–10. issn: 20411723. (2020) (Dec. 2020).

22. Smith, I. et al. Evaluation of RNAi and CRISPR technologies by large-scale gene expression profiling in the Connectivity Map. PLoS Biology 15 (ed Freeman, T.) Publisher: Public Library of Science, e2003213. issn: 15457885. http://dx.plos.org/10.1371/journal.pbio.2003213 (2018) (Nov. 2017).

23. Zhu, J., et al. Prediction of drug efficacy from transcriptional profiles with deep learning. Nature Biotechnology. Publisher: Nature Publishing Group, 1–9. issn: 15461696. http://www.nature.com/articles/s41587-021-00946-z (2021) (June 2021).

24. Cimini, B. A., et al. Optimizing the Cell Painting assay for image-based profiling. en. Nature Protocols 18. Number: 7 Publisher: Nature Publishing Group, 1981–2013. issn: 1750-2799. https://www.nature.com/articles/s41596-023-00840-9 (2023) (July 2023).

25. Lundholt, B. K., Scudder, K. M. & Pagliaro, L. A Simple Technique for Reducing Edge Effect in Cell-Based Assays. en. Journal of Biomolecular Screening 8. Publisher: SAGE Publications Inc STM, 566–570. issn: 1087-0571. 10.1177/1087057103256465 (2023) (Oct. 2003).

26. Gerry, C. J. & Schreiber, S. L. Recent achievements and current trajectories of diversity-oriented synthesis. Current Opinion in Chemical Biology. Next Generation Therapeutics 56, 1–9. issn: 1367-5931. https://www.sciencedirect.com/science/article/pii/S1367593119301000(2023) (June 2020).

27. Enache, O. M. et al. The GCTx format and cmap*{*Py, R, M, J*}* packages: resources for optimized storage and integrated traversal of annotated dense matrices. Bioinformatics 35 (ed Kelso, J.) Publisher: Narnia, 1427–1429. issn: 1367-4803. https://academic.oup.com/bioinformatics/article/35/8/1427/5094509 (2019) (Apr. 2019).

